# NON-EQUILIBRIUM BELOUSOV–ZHABOTINSKY REACTION DESCRIBES THE SPONTANEOUS ACTIVITY OF THE BRAIN

**DOI:** 10.1101/863324

**Authors:** Y-idi Zhang, Shan Guo, Mingzhu Sun, Arturo Tozzi, Xin Zhao

**Affiliations:** Institute of Robotics and Automatic Information System, Nankai University, Tianjin, China; Center for Nonlinear Science, Department of Physics, University of North Texas, Denton, Texas 76203, USA, 1155 Union Circle, #311427

**Keywords:** central nervous system, chaos, chemical reaction, spontaneous activity, BOLD activity, nonlinear dynamics

## Abstract

Belousov–Zhabotinsky (BZ) reactions describe chemical patterns in terms of non-equilibrium thermodynamics, chaotic evolution of nonlinear oscillators, excitability under the influence of peculiar chemical stimulations. In touch with this chemical model, we hypothesized that the nonlinear brain electric activity at the edge of the chaos could be triggered by neural oscillation equipped with BZ dynamics, and that changes in neural activity might be correlated with the transient occurrence of (either short and long-range) BZ-like reactions in cortical subareas. To prove our hypothesis of neuronal waves driven by BZ-like processes, we evaluated fMRI movies that assess in vivo BOLD resting state activity of the human brain. We found that the spontaneous activity of the brain display features fully overlapping the recently-introduced local circuits based on BZ chemical reaction. Therefore, neuronal paths during spontaneous activity of the brain match BZ dynamics’ previsions. Our results point towards the brain as crossed by diffusive nonlinear neural oscillations patterns that are predictable. Furthermore, our results suggest that chaotic dynamics arise from nothing else than the network arrangements subtending physical and biological systems.

## INTRODUCTION

Belousov–Zhabotinsky (BZ) reactions describe a nonlinear chemical oscillator in the framework of non-equilibrium thermodynamics. In presence of bromine and an acid, a chemical reaction is produced that is far from equilibrium and evolves chaotically. This intriguing chemical model of noise-induced order has been used for chemical computations that, unlike conventional computing, rely on geometrically constrained excitable chemical medium and use changes in BZ reagents’ concentrations to transmit information. This approach shaped structured excitable media that have been proven useful to describe the features of different physical and biological systems, such as, e.g., image processing, Voronoi diagram, logical computation. Starting from BZ reaction, Zhang et al. (2012), designed a planar geometrical binary adder chemical device that realizes a two-bit, or even multi-bit logical computation. Sun and Zhao’s (2013) simulations showed how one-bit decoders can be extended through cascade methods to design two-bit, three-bit, or higher bit binary decoders. Therefore, chemical realization of decoders can drive the building of more sophisticated functions based on BZ reaction.

Here we tackle the issue from the standpoint of the brain cortical activity. Indeed, brain oscillations have common ground with BZ reactions: both display chaotic features and self-organizing activity under the influence of specific stimuli; both exhibit “excitability”, i.e., the occurrence of patterns developing in apparently quiescent media. Given these premises and points in common, we hypothesize that BZ mechanisms might explain the oscillatory behavior of fMRI BOLD activation during different cognitive activities and/or tasks and, in particular, during spontaneous activity of the brain. Our aim is to introduce a model of nervous activity where oscillations arising from single areas or subareas diffuse-inhomogeneously and unergodically-towards different cortical locations. We provide the required BZ equations and compare theoretical results from BZ simulations with available real neurodata in subjects during mind wandering ans spontaneous activity of the brain. In sum, starting from the activity density in different areas of the brain, we aim to prove that oscillation migration follows BZ dynamics in a way that is predictable. We show how the diffusion of the neural waves - towards both nearby and far apart areas - can be described through BZ dynamics equipped with random oscillation propagation. The matching of theoretical BZ models and real patterns of neural activity aims to achieve three main goals: 1) to demonstrate that the brain is activated through a recognizable diffusion pattern of nervous structures and wave spread; 2) to provide neuroscientists with a significant information: the possibility to reproduce and standardize the propagation of neural waves in different cognitive activities; 3) to show how the chaotic, non-linear activity occurring during brain activity is not correlated with specific biochemical reactions, rather it requires nothing else that a peculiar structure of the subtending neural networks. Therefore, chaos in the brain comes just from the constrained shape of circuits, nodes and edges.

## MATERIALS AND METHODS

Our aim is to correlate the simulations introduced in previous papers with brain activities. In order to pursue our task, we compared neurodata from real fMRI movies with the recently-developed Multi-bit binary decoder based on Belousov-Zhabotinsky reaction.

### fMRI movies

Structures of high dimensionality, termed “lag threads”, can be found in the brain during spontaneous activity (Mitra et al., 2015). Lag threads consist of multiple, highly reproducible temporal sequences. We retrospectively evaluated free-available published video frames showing lag threads computed from real BOLD resting state rs-fMRI data. The group consisted of 688 subjects from the Harvard-MGH Brain Genomics Superstruct Project. We examined four sets of movies including sagittal, trasversal, and coronal sections for a total of 54 Images, taken from the videos: http://www.pnas.org/content/suppl/2015/03/24/1503960112.DCSupplemental (Threads 1, 2, 3 and 4). Therefore, according to Mitra et al.’s (2015) claims, brain activities encompass single or multiple lag threads. In the sequel, we will provide an effort to describe lag threads in terms of BZ reactions.

### Multi-bit binary decoder based on Belousov-Zhabotinsky reaction: experimental design

In previous studies, BZ reactions have been described in terms of chemical calculations with simple (One-way propagation, Osmotic propagation, Delayed propagation) and complex structures (Adders, Memory, Decoders). For further details, see: Zhang et al. (2012); Sun and Zhao (2013); Guo et al. (2014). The binary adder unit was built with simple straight line boundaries, by integrating existing functional structures and their implementations, such as T-shaped structures, unidirectional transmission structures, and cross-propagation structures. The unit performs the addition of binary information via the geometrically constrained structure, with no need of ruling clocks or parameters changes during the process. These single-bit binary adder units were then coupled to produce a two-bit binary adder, using a method similar to the construction of adders in digital circuits. With the same strategy, higher bit binary adders can be implemented, so that the structures of multi-bit binary decoders become complex as number of bits increases.

The authors designed multibit decoders using a cascade method, that linked n-bit decoders (n ≥ 2) with (n − 1)-bit decoders, so that a n-bit decoder could be built based on a (n − 1)-bit decoder. Each n-bit decoder includes the simpler structure of a (n − 1)-bit decoder. Simulations results demonstrated that this type of circuit achieves decoding functions. The authors performed the whole simulations towards the realization of a combinational logic circuit, i.e., a multi-bit binary decoder that converts binary information from *n* input lines to a maximum of 2*n* unique output lines.

### Correspondences between videos and simulations

Once established that an available circuit is able to describe and simulate the functionality of BZ reactions, we implemented its design to assess both simple and more intricate neural patterns related to spontaneous activity of the brain. To achieve our goal, we started from the above-mentioned claim by Mitra et al. (2015) that a brain activity encompasses single or multiple lag threads. We used the video frames of BOLD resting state rs-fMRI activity as the domain of BZ reaction, looking for the correspondence between circuit simulations and spontaneous activity of the brain. In other words, we tried to assess and reproduce the occurrence of possible BZ reaction in the available video frames. Once set the location of signals according to the videos, we simulated active signals in the corresponding brain areas. The BZ model’s features of propagation suggest that: a) single signals can span the narrow gap; b) single signals cannot cross the wide gap; c) when the signals meet, they annihilate; d) a signal transmission occurs along the channels (Sun and Zhao, 2013). In terms of BZ networks, it may be postulated that each thread encompasses two elements: nodes and nodes propagation sequence. The governing ideas are: 1. Node; 2. Propagation sequence between nodes; 3. Single lag thread; 4. Multiple lag threads. Therefore, to simulate brain activity, it is mandatory to simulate both a single lag thread and a superposition of multiple lagging threads. To build the model as simple as possible, the nodes stand here for single brain subareas equipped with input and output channels (**Figure 1.1**). To achieve the required propagation sequence between nodes, we used one-way transmission structures (**Figure 1.2**). The combination of nodes and propagation sequence generates at first single lag threads (**Figure 1.3**), then multiple lag threads (see **Figure 1.4**, that illustrates the superposition of Thread 1 and Thread 2).

**Figure 1.**
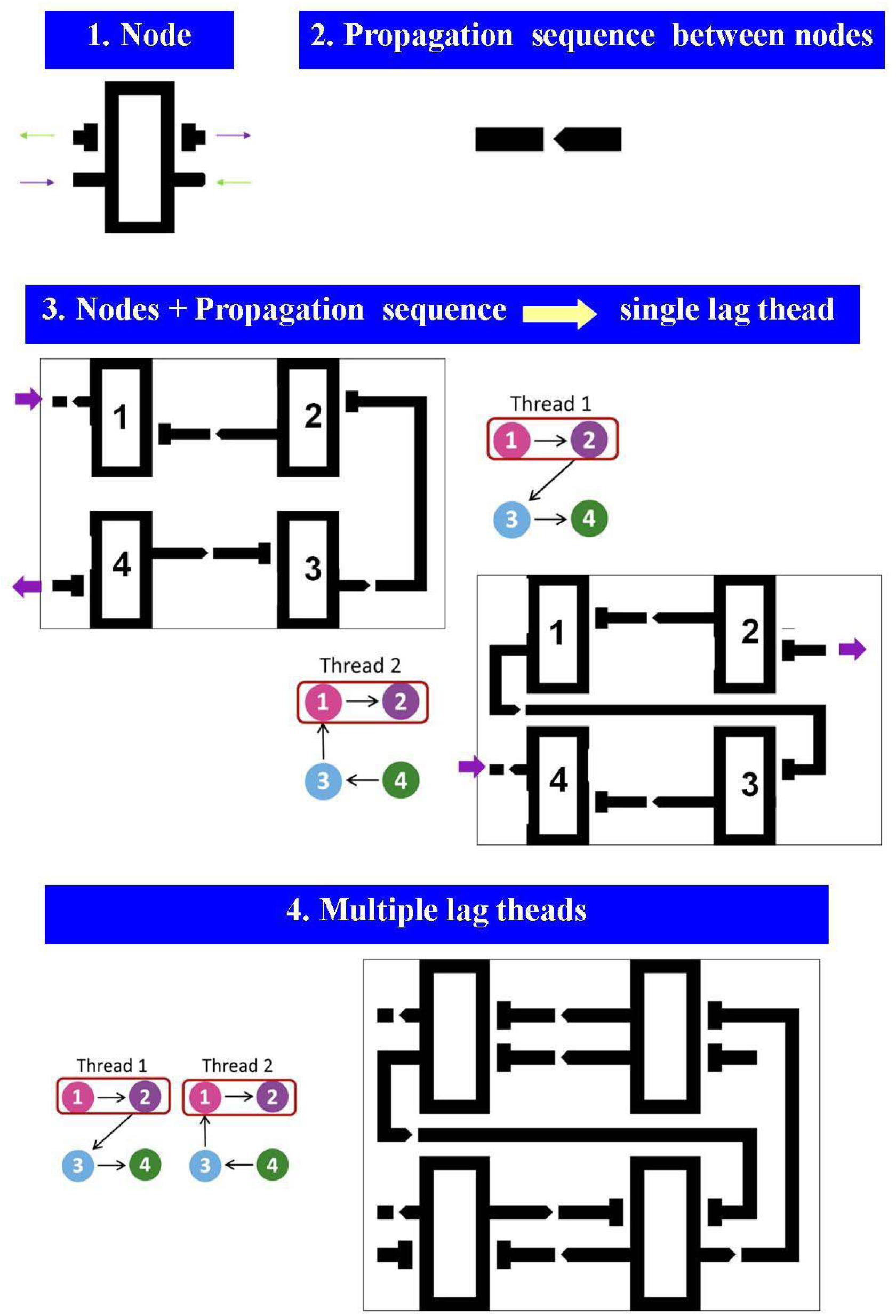
Spontaneous activity of the brain described in terms of the Multi-bit binary decoder based on Belousov-Zhabotinsky reaction. Note that circuits that illustrate and simulate multiple lag threads structures can be achieved starting from a very few basic components. See text for further details.

## RESULTS

The diffusive paths and changes in BOLD neuronal activity can be described in terms of BZ models and their master equations. Indeed, phenomena and dynamics correlated with BZ reaction where found in the video frames of BOLD resting state rs-fMRI activity.

### Correspondence between video and appearance simulation

As stated in the previous paragraphs, BZ-based frameworks predict that: a) single signals can span the narrow gap; b) single signals cannot cross the wide gap; c) when signals meet, they annihilate; d) signal transmission occurs along the channels. We found that at least some of these predictions are confirmed in the neurodata, because matching features occur between the brain signals detectable in the real movies and the simulations performed through BZ-based circuits. Indeed, we found that single signals span the narrow gaps. This happens almost everywhere, in every time window. For example, it occurs from 0.8s to 1.1s in the video of thread 1 (**Figure 2, upper part**). This phenomenon takes place in both real videos and BZ simulations. We also found that, when the signals meet, they annihilate. For example, it occurs from 0.7s to 1.1s in the video of thread 4 (**Figure 2, lower part**). This phenomenon takes place both in video and BZ simulation, in many brain areas. Concerning the signal transmission along the channel, this is an event that occurs almost everywhere in the signal transmission throughout the whole brain. In particular, we noticed that signals favor to span the narrow gap, rather than going along the channel.

**Figure 2.**
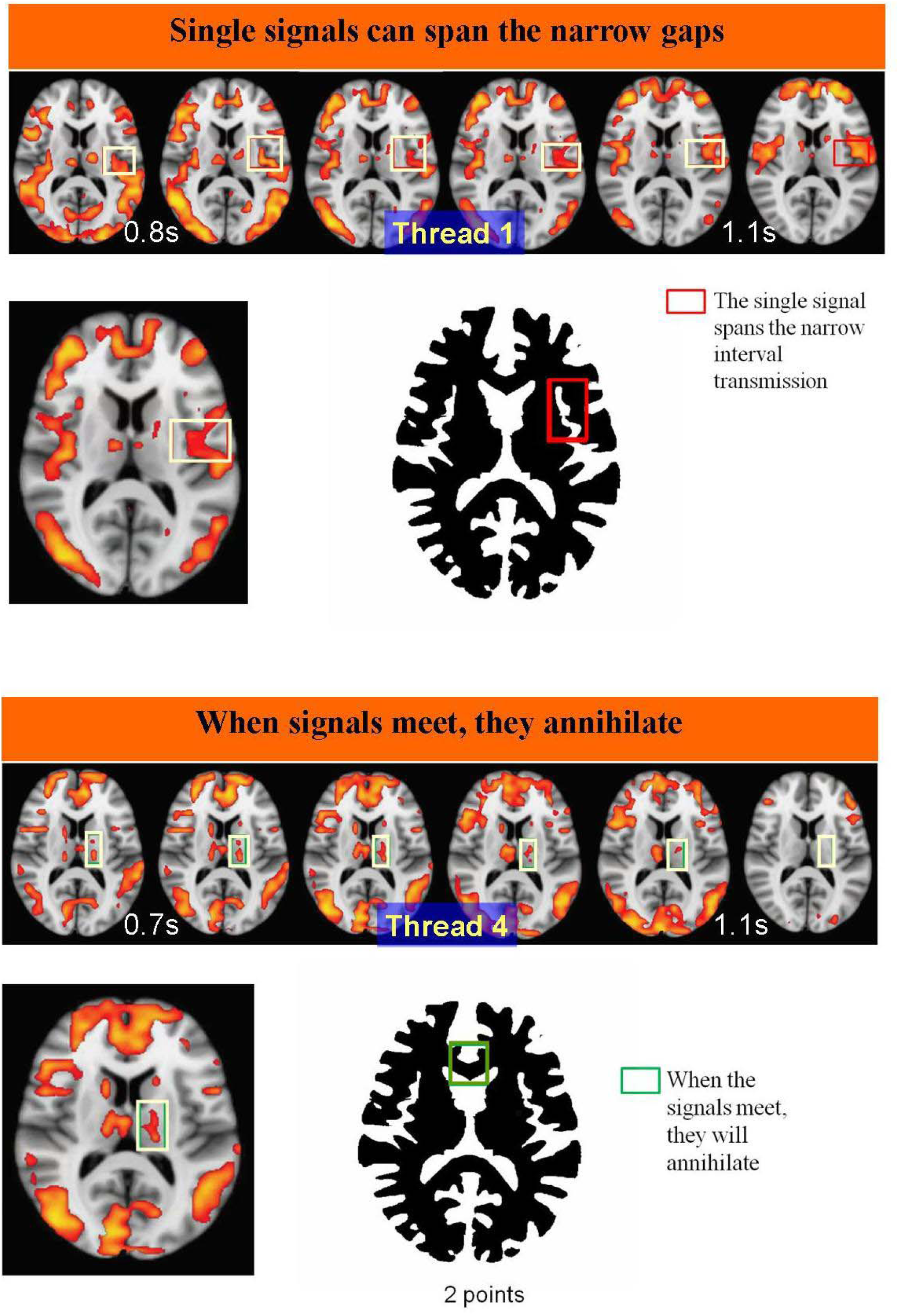
Correspondence between lag thread frames and BZ simulation permits the correlation between lag thread frames and BZ circuit. See text for further details.

### Building lag threads

When comparing the theoretical curves from RD simulations with the ones extracted from real data, we noticed remarkable superimposition between the two paths. We took, for example, two random lag threads propagating through a few nodes and performed the BZ reaction to describe both single and multiple lag threads. **Figure 3 (upper and middle parts)**, based on the video available by Mitra, illustrates how to build the threads 1 and 2 and how to connect them. Note that the two lag structures can be used also to control the lag time. Indeed, due to the scarce homogeneity of the connections between nodes, it takes different time for signals to pass through one node to another. Since the signals are transmitted throughout the channels at the same speed, lag time can be monitored and modified in two ways: a) by changing the channel length between nodes; b) in addition, since the speed of the signal will slow down as it passes through the gap, lag time can be changed by adding novel channels gaps (see **Figure 3, lower part**).

**Figure 3:**
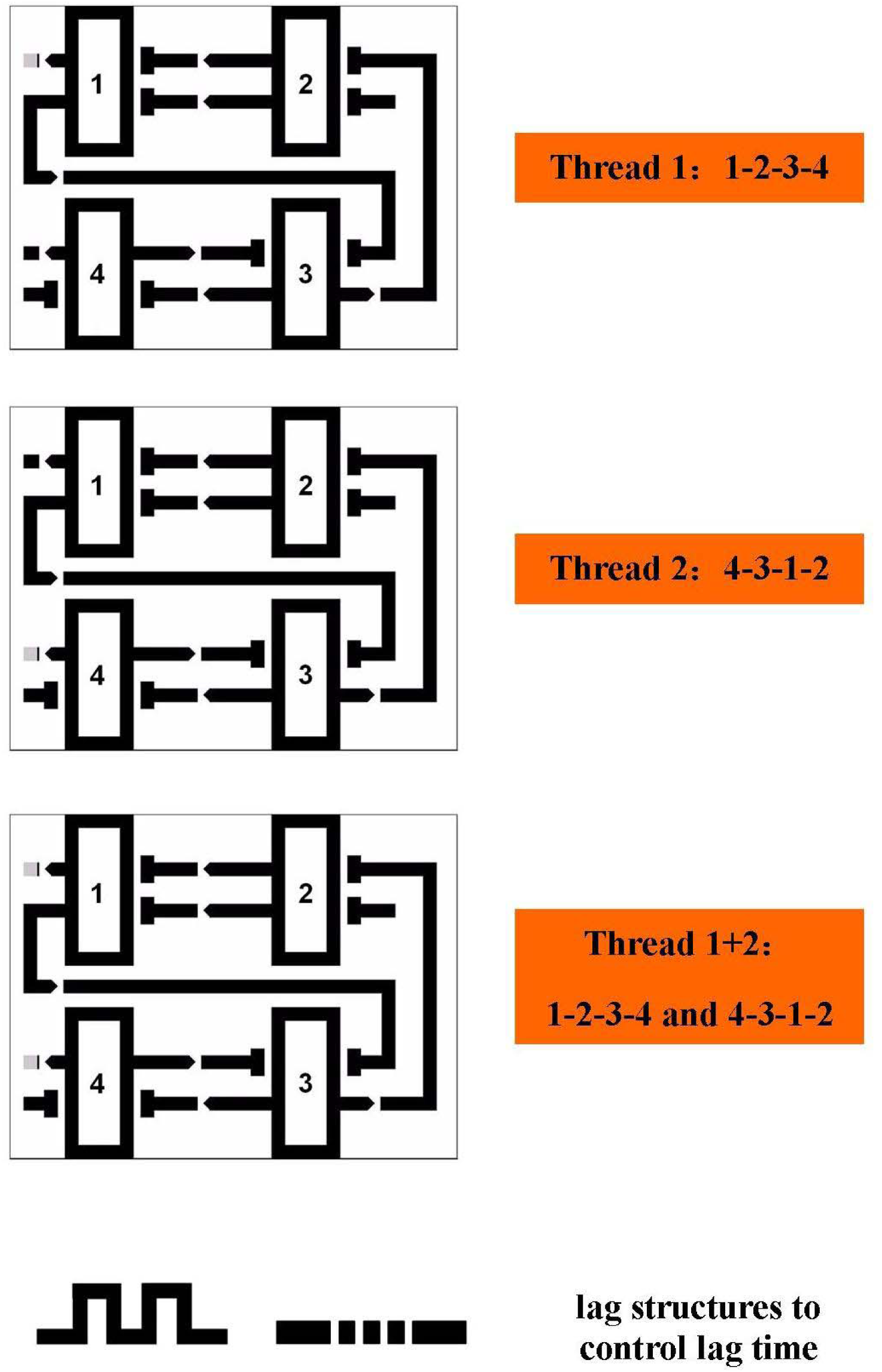
Simulation of BZ circuits, starting from the parameters described in video frames of BOLD resting state rs-fMRI activity. The Figure suggests that interactions between brain areas are quantifiable in terms of BZ equations.

## DISCUSSION

We started from a recently-developed chemical signal processor based on the BZ reaction platform, that uses single-bit binary adder units and a two-bit binary adder. We found that the pattern of neuronal activity predicted by BZ models’ simulations fully matches the findings detected in the brain of subjects during spontaneous activity. Or, in other words, the behavior of the BZ reaction is very similar to the behavior of brain functions. This phenomenon suggests that the use of BZ reactions to simulate brain activity is promising. Indeed, the design of these logical devices, based on the space-time interaction of travelling excitation waves, is well-suited for experimental implementation in neuroscience (see also: Gomez-Molina et al., 2017).

### Next to come

Concerning the limitations of this study, a series of constraints can be drawn. Because of the intricate nature of excitable medium and the high dependency on synchronized inputs, careful control of the input signals is required to realize the binary addition, both in chemical experiments and in brain dynamics. In this papers, we stated that nodes stand for brain subareas equipped with inputs and outputs. This is because we choose a coarse-grained macro-level of observation and analysis. In turn, if we consider a coarse-grained micro-level of observation and analysis, nodes might also stand for single neurons. Therefore, a few crucial questions arise: is it possible to simplify nodes, without providing a sharp description of their specificity and functionality? The answer is positive, because simplifying nodes does not affect the description of threads. Indeed, BZ reaction has been able to provide a very good description of single threads and their subsequent superposition. Further, is it feasible to achieve a multi-threaded superposition? In this paper, we implemented just a two-threads superposition, while a more complicated superposition of multiple threads would require further exploration and more specific examples: in particular, we would need to take into account not just the case of spontaneous activity of the brain, but also the case of brain activities following specific tasks. When examining the very concept of lag thread, the term “lag” is a key innovation point, suggesting that the concept of lag into needs to be incorporated in the circuit. However, here we ask: is it feasible to weight threads? The suggestion of Mitra et al. (2015) - that different threads occupy different weights in brain activities - is difficult to achieve and describe in the context of BZ reaction. Our theoretical idea was to design a structure able to treat signals from different threads differently. For example, suppose that the weight of thread 1 is higher than the weight of thread When the signals in thread 1 and 2 cross simultaneously the structure, signal 1 will pass and destroy signal 2: this means that a weight difference between different threads takes place. In turn, if the sole signal 2 crosses the circuit, it will not be affected. We conjecture that how (and how many) these circuit structures are implemented will stand for a central method for adjusting thread weights.

### Assessing chaos through networks arrangement

A BZ model for neural activity predicts that the subtle balance between oscillations gives rise to chaotic patterns. In touch with BZ reactions, the brain has been described as a complex, non-linear system operating at the edge of chaos, with inter-dependent components exhibiting spontaneous self-organization/emergent properties (Tognoli and Kelso, 2013; Fraiman and Chialvo, 2012; Zare and Grigolini, 2013; Xu and Wang, 2014). The brain is a phase space in which particle movements take place, with different trajectories displaying different paths (Watanabe et al., 2013; Yan et al., 2013; Kim and Lim 2015; Wang et al., 2017). It has been suggested that the brain phase space displays funnel-like locations where trajectories converge towards the shortest path as time progresses (Tozzi et al., 2016; Sengupta et al., 2016). Other neuroscientists suggested that brain function does not exhibit erratic brain dynamics nor attractors, rather a stable sequence of transient heteroclinic channels (Afraimovich et al., 2013). Further, concepts such as communication-through-coherence (Deco and Jirsa, 2012) and collective movements (Touboul 2012; Tozzi, 2015) must be taken into account. Summarizing, different nonlinear functional regimes occurring in the brain phase space have been described, both in central nervous systems and in artificial neural networks (Tozzi et al., 2016). In touch with brain issues, the solutions of BZ equations describe a wide range of nonlinear behaviors, including the formation of travelling waves and wave-like phenomena, as well as other self-organized patterns. The network models and circuits built by the above-described Authors display several logical devices, such as Boolean logic gates, adders, counters, memory cells. This means that clever geometrical arrangements of channels for excitation wave propagation are able to describe nonlinear dynamics. Also this suggests that the key to understand chaotic dynamics lies inside the structure and arrangement of physical and biological networks.

## REFERENCES

1) V. Afraimovich, I. Tristan, P. Varona, M. Rabinovich. 2013. Transient Dynamics in Complex Systems: Heteroclinic Sequences with Multidimensional Unstable Manifolds. Discontinuity, Nonlinearity and Complexity 2(1): 21–41.

2) Deca D. 2017. Grid Cells-From Data Acquisition to Hardware Implementation: A Model for Connectome-Oriented Neuroscience. In: The Physics of the Mind and Brain Disorders: Integrated Neural Circuits Supporting the Emergence of Mind, edited by Opris J and Casanova MF. New York, Springer; Series in Cognitive and Neural Systems. Pages 493-511. ISBN: 978-3-319-29674-6. DOI 10.1007/978-3-319-29674-6_9.

3) Deco G, Jirsa VK. 2012 Ongoing cortical activity at rest: criticality, multistability, and ghost attractors. J Neurosci. 7;32(10):3366–75. doi: 10.1523/JNEUROSCI.2523-11.2012.

4) Fraiman D, Chialvo DR. 2012. What kind of noise is brain noise: Anomalous scaling behavior of the resting brain activity fluctuations. Frontiers in Physiology, 3 JUL(July), 1–11. http://doi.org/10.3389/fphys.2012.00307

5) Gomez-Molina JF, Corredor M, Restrepo-Velasquez AA, Ricoy UM. 2017. Computer models for ions under electric and magnetic fields: random walks and relocation of calcium in dendrites depends on timing and population type. IFMBE proceedings. In: VII Latin American Congress on Biomedical Engineering CLAIB 2016, Bucaramanga, Santander, Colombia, October 26th -28th, 2016. DOI: 10.1007/978-981-10-4086-3_175

6) Guo S, Sun M-Z,Han J-D, Zhao X. 2014. Digital Comparator in Excitable Chemical media. Int. Journ. of Unconventional Computing, Vol. 0, pp. 1–15, 2014.

7) Kim SY, Lim W. 2015. Frequency-domain order parameters for the burst and spike synchronization transitions of bursting neurons. CognNeurodyn. 2015 Aug;9(4):411–21. doi: 10.1007/s11571-015-9334-4.

8) Mitra A, Synder AZ, Blazey T, Raichle ME (2015). Lag threads organize the brain’s intrinsic activity. Proc Natl Acad Sci USA 112(17):2235–2244.

9) Xu X, Wang R. 2014. Neurodynamics of up and down transitions in a single neuron. CognNeurodyn. 8(6):509–15. doi: 10.1007/s11571-014-9298-9.

10) Sengupta B, Tozzi A, Cooray GK, Douglas PK, Friston KJ. 2016. Towards a Neuronal Gauge Theory. PLOS Biology, 14(3), e1002400. http://doi.org/10.1371/journal.pbio.1002400

11) Sun M-Z and Zhao X. 2013. Multi-bit binary decoder based on Belousov-Zhabotinsky reaction. The Journal of Chemical Physics 138, 114106 (2013); doi: 10.1063/1.4794995

12) Tognoli E, Kelso JS. 2013. On the Brain’s Dynamical Complexity: Coupling and Causal Influences Across Spatiotemporal Scales. Advances in Cognitive Neurodynamics (III), (Iii), 259–265. http://doi.org/10.1007/978-94-007-4792-0.

13) Touboul J. 2012. Mean-field equations for stochastic firing-rate neural fields with delays: Derivation and noise-induced transitions. Physica D: Nonlinear Phenomena. 241 (15):1223–1244. doi:10.1016/j.physd.2012.03.010

14) Tozzi A, Fla T, Peters PJ. 2016. Building a minimum frustration framework for brain functions in long timescales. J Neurosci Res. DOI: 10.1002/jnr.23748

15) Tozzi A. 2015 Information processing in the CNS: a supramolecular chemistry? CognNeurodyn. 2015 Oct;9(5):463–77. doi: 10.1007/s11571-015-9337-1. Review.

16) Wang Y, Wang R, Zhu Y. 2017. Optimal path-finding through mental exploration based on neural energy field gradients. CognNeurodyn. 11(1):99–111. doi: 10.1007/s11571-016-9412-2.

17) Watanabe, T., Hirose, S., Wada, H., Imai, Y., Machida, T., Shirouzu, I., Masuda, N. 2013. A pairwise maximum entropy model accurately describes resting-state human brain networks. Nature Communications, 4, 1370. http://doi.org/10.1038/ncomms2388

18) Zare M, Grigolini P. 2013. Chaos, Solitons & Fractals Criticality and avalanches in neural networks. Chaos, Solitons and Fractals: The Interdisciplinary Journal of Nonlinear Science, and Nonequilibrium and Complex Phenomena, 55, 80–94. http://doi.org/10.1016/j.chaos.2013.05.009.

19) Zhang G-M, Wong I, Chou M-T, Zhao X. 2012. Towards constructing multi-bit binary adder based on Belousov-Zhabotinsky reaction. Citation: J. Chem. Phys. 136, 164108 (2012); doi: 10.1063/1.3702846.

